# Molecular characterization of *Chlamydomonas reinhardtii* telomeres and telomerase mutants

**DOI:** 10.1101/519991

**Authors:** Stephan Eberhard, Sona Valuchova, Julie Ravat, Pascale Jolivet, Sandrine Bujaldon, Stéphane D. Lemaire, Francis-André Wollman, Maria Teresa Teixeira, Karel Riha, Zhou Xu

## Abstract

Telomeres are repeated sequences found at the end of the linear chromosomes of most eukaryotes and are required for chromosome integrity. They shorten with each cell division because of the end-replication problem. Expression of the reverse transcriptase telomerase allows for extension of telomeric repeats to counteract telomere shortening. Although *Chlamydomonas reinhardtii*, a photosynthetic unicellular green alga, is widely used as a model organism in photosynthesis and flagella research, and for biotechnological applications, the biology of its telomeres has not been investigated in depth. Here, we show that the *C. reinhardtii* (TTTTAGGG)_n_ telomeric repeats are mostly non-degenerate and that the telomeres form a protective structure, ending with a 3′ overhang. While telomere size and length distributions are stable under various standard growth conditions, they vary substantially between 12 genetically close reference strains. Finally, we identify *CrTERT*, the gene encoding the catalytic subunit of telomerase and show that mutants of this gene display an “ever shortening telomere” phenotype and eventually enter replicative senescence, demonstrating that telomerase is required for long-term maintenance of telomeres in *C. reinhardtii*.

## INTRODUCTION

Photosynthetic algae are in the highlight of basic and applied research, not only because of their core role for Earth’s biosphere in oxygen evolution and carbon fixation, but also because of their increased use in biotechnology for the production of proteins, bulk chemicals and high-value molecules (Scaife et al., 2015; Scranton et al., 2015). Thus, a detailed understanding of algal physiology, including their cell cycle, cell growth and genome integrity, is of critical importance. *Chlamydomonas reinhardtii*, also referred to as the “photosynthetic yeast” (Rochaix, 1995), is the most prominent model organism in the green algae lineage and is widely used for biotechnological applications as well as to study fundamental processes, such as photosynthesis and cilia structure and function (Harris, 2001; Sasso et al., 2018).

In eukaryotes, telomeres are repeated sequences found at the extremities of linear chromosomes. They are important for chromosome integrity and may limit cell proliferation capacity in some organisms. By progressively shortening with each cell cycle because of the end-replication problem, telomeres eventually become too short and trigger a cell cycle arrest termed replicative senescence (Harley et al., 1990; Lundblad and Szostak, 1989). Most unicellular eukaryotes and germ, stem and cancer cells in multicellular organisms, counteract telomere shortening by expressing telomerase, an enzyme that adds *de novo* telomere sequences and allows for an unlimited proliferation potential (Pfeiffer and Lingner, 2013; Wu et al., 2017). Despite the crucial functions of telomeres and telomerase in maintaining genome stability and controlling cell proliferation in many model organisms including plants, ciliates, fungi and mammals (Fulcher et al., 2014), telomere biology in algae remains to be investigated in depth.

To our knowledge, only a handful of studies on *C. reinhardtii* telomeres have been published. Early studies published in the 90s showed that: (i) *C. reinhardtii* telomeres are composed of TTTTAGGG repeats, which are different from the *Arabidopsis*-type TTTAGGG sequence (Petracek et al., 1990); (ii) the size of cloned telomeric repeats ranges from 300 to 600 bp (Hails et al., 1995; Petracek et al., 1990); (iii) they form G-quadruplex structures *in vitro* (Petracek and Berman, 1992), and (iv) the Gbp1 protein potentially binds to telomeres (Johnston et al., 1999; Petracek et al., 1994). More recently, bioinformatic studies focused on the evolutionary relationships of telomere sequences in green algae (Fulneckova et al., 2012; Fulneckova et al., 2015). Finally, a broad study of telomerase activity in green algae revealed that telomerase activity in *C. reinhardtii* extracts is low or not detectable (Fulneckova et al., 2013).

To gain a better understanding of *C. reinhardtii* telomere structure and maintenance, we investigated telomere sequence and end structure, analyzed telomere length distribution across different reference strains, identified *CrTERT*, the gene encoding the catalytic subunit of telomerase, and provided a genetic analysis of telomerase function, thus opening new avenues of research on telomere dynamics, proliferation potential and genome integrity in *C. reinhardtii*.

## RESULTS

### *C. reinhardtii* telomeric repeats are mostly non-degenerate with few low-frequency variants

In their seminal paper, Petracek *et al*. cloned and sequenced a limited number of *C. reinhardtii* telomeric repeats, revealing their canonical TTTTAGGG sequence (Petracek et al., 1990). Telomeric repeats are also identifiable in 18 out of 34 chromosome ends on the available v5.5 genome sequence of *C. reinhardtii* (https://phytozome.jgi.doe.gov; **Supplemental Figure S1A**). As the sequenced genome shows some telomeric repeat variations, we analyzed telomeric repeat sequences on a larger scale and looked for putative variants of the canonical telomere sequence. We amplified telomeres by a PCR-based method (Forstemann et al., 2000) using a forward primer specific to a conserved subtelomere-telomere junction common to 10 telomeres from 8 different chromosomes (**Supplemental Figure S1A and S1B**). The reverse primer was universal and annealed to a sequence of cytosines, artificially added at the 3’-end of the telomeres by terminal transferase reaction. After cloning into a plasmid and sequencing, we analyzed 32 telomere sequences, encompassing 709 repeats. We found that ~90% (n = 636) of the repeats corresponded to the canonical sequence TTTTAGGG. We also detected variants such as TTTAGGG (corresponding to the canonical *A. thaliana* sequence, n = 37, either at the subtelomere-telomere junction, n = 24, or elsewhere, n = 13) or TTTTTAGGG (n = 13) and TTTTGGG (n = 8) (**Table 1** and **Supplemental Figure S1B**). These three variants were found in at least two independent clones at the same position in the telomere sequence, thus likely representing true low-frequency variants and not sequencing errors. We also detected sequence variants that occurred only in single clones (n = 15) and for which PCR and/or sequencing errors can therefore not be ruled out. We conclude that *C. reinhardtii* telomeric repeats are mostly non-degenerate with few low-frequency variants.

**Table 1:** Frequency of telomeric repeats motifs determined by telomere PCR and sequencing of 32 independent clones

| Sequence | n | freq. |
| --- | --- | --- |
| TTTTAGGG | 636 | 89.7% |
| TTTAGGG | 37 | 5.2% |
| TTTTTAGGG | 13 | 1.8% |
| TTTTGGG | 8 | 1.1% |
| Others | 15 | 2.1% |

### *C. reinhardtii* telomeres form a non-nucleosomal protective structure, bear a 3′ overhang and show no evidence for blunt ends

The protective structure formed by telomeric DNA bound by specific proteins is critical for telomere functions (Palm and de Lange, 2008). To test the presence of such a structure at *C. reinhardtii* telomeres, we performed a micrococcal nuclease (MNase) digestion of nuclei and asked whether telomere DNA would be protected from its activity. When nuclei were subjected to increasing amounts of MNase, nucleosomal DNA was protected from digestion and migrated at ~150 bp based on ethidium bromide staining (**Figure 1A**, left), as expected (Clark, 2010). Intermediate digestion products migrated in a typical ladder pattern corresponding to di-nucleosomes, tri-nucleosomes and higher order structures. Strikingly, Southern blotting with a radioactive telomeric probe revealed that telomeric DNA was protected from MNase digestion in a non-nucleosomal pattern (**Figure 1A** right). As a control, the same membrane was stripped and probed for 18S rDNA, revealing the canonical nucleosome structure (**Figure 1A**, middle). The size of the protected telomeric DNA was in the range of 200-700 bp, which could correspond to the full telomere length. This result suggests that telomeric DNA might be fully associated with and protected by protein complexes in a non-nucleosomal structure, similar to telosomes as observed in yeast for example (Wright et al., 1992).

**Figure 1:**
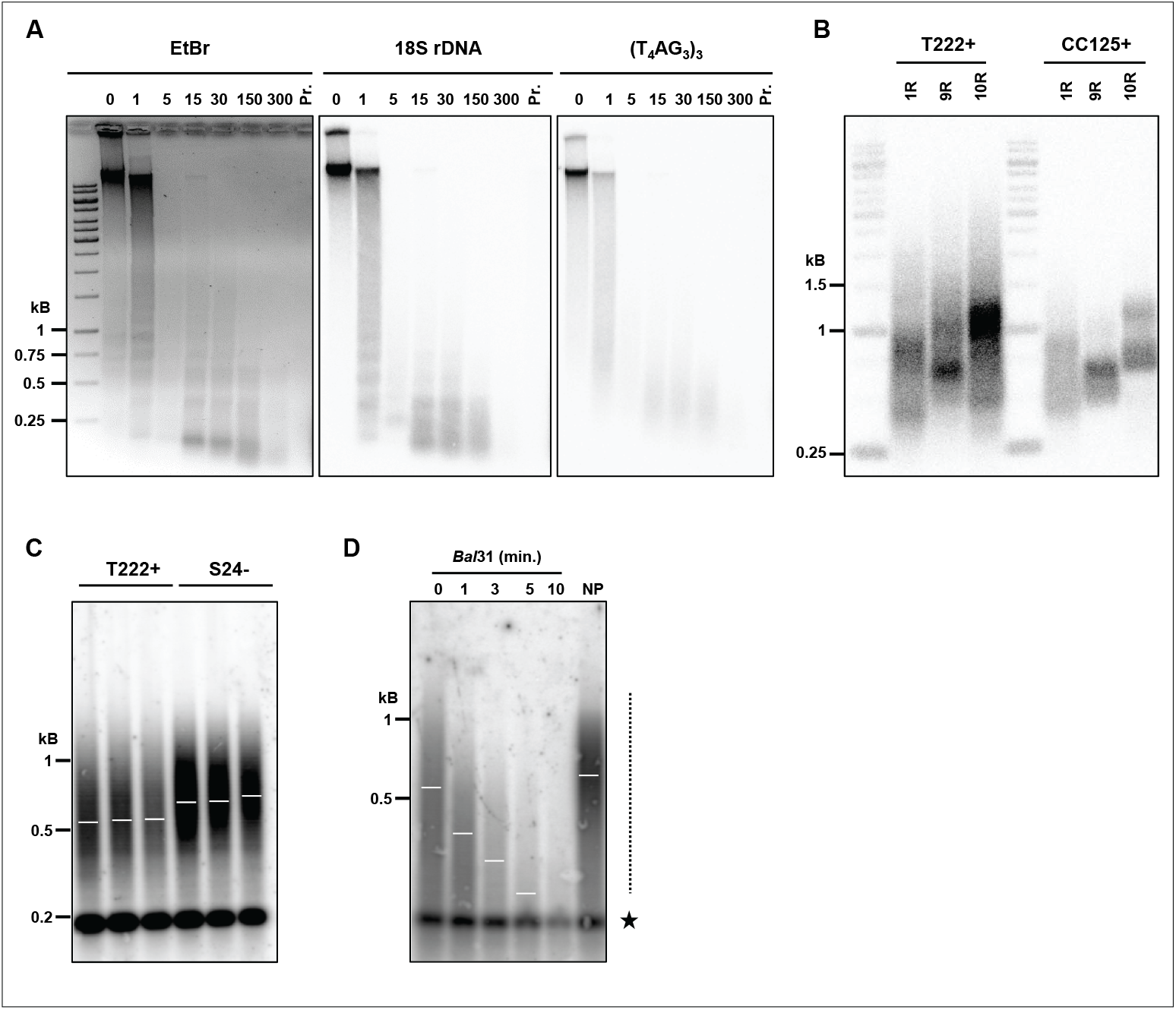
Structural characterization of *C. reinhardtii* telomeres. **(A)** Characterization of *C. reinhardtii* telosomes by MNase digestion of gDNA (left panel; “EtBr”: ethidium bromide staining of the migration gel) and Southern analysis with a telomeric specific probe (right panel; (T_4_AG_3_)_3_: radiolabeled probe). The membrane was then stripped and probed again with an 18S rDNA probe (middle panel). **(B)** PETRA was used to amplify specific telomeres from strains T222+ and CC125+, and analyzed by Southern blotting using the telomere-specific probe (TTTTAGGG)_3_ (see also **Supplemental Figure S1C**). **(C)** T222+ and S24-strains were subcloned and three subclones were independently grown in liquid cultures until stationary phase and subsequently analyzed by TRF Southern blot. **(D)** Genomic DNA was subjected to *Bal*31 digestion for 1 to 10 minutes. Digested products were column-purified and then digested with the restriction enzyme cocktail, separated by electrophoresis and analyzed by TRF Southern blot. 0: no *Bal*31 digestion, but gDNA was column-purified before digestion by the restriction enzymes. NP: gDNA was directly analyzed by TRF Southern blot, with No column-Purification. Dashed line: smear corresponding to telomeres. Star: *Bal*31-insensitive band, corresponding to interstitial telomeric repeat (see also **Supplemental Figure S1F**).

The chromosome end-structure determines the protection strategies employed to cap the telomere. In many species, telomeres end with a 5′ to 3′ single-stranded overhang, important for the protective t-loop structure in human telomeres, telomerase recruitment and binding of specific proteins, such as the CST and Ku complexes (Giraud-Panis et al., 2010; Palm and de Lange, 2008; Wellinger and Zakian, 2012). As it was reported that the Gbp1 protein preferentially binds single-stranded *C. reinhardtii* telomeric DNA (Johnston et al., 1999), the presence of a 3′ overhang would be consistent with a role of Gbp1 at telomeres. In order to experimentally test the presence of a 3′ overhang at *C. reinhardtii* telomeres, we performed <u>P</u>rimer <u>E</u>xtension <u>T</u>elomere <u>R</u>epeat <u>A</u>mplification (PETRA) (Heacock et al., 2004)). PETRA requires the annealing of an adaptor primer (PETRA-T) to the overhang. After primer extension, the telomere was PCR-amplified using a unique subtelomeric forward primer and a reverse primer (PETRA-A) complementary to a tag sequence present in PETRA-T (**Supplemental Figure S1C**). Successful amplification by PETRA is indicative of the presence of a 3′ overhang. Using primers specific for three different telomeres (1R, 9R and 10R), we found robust amplification of PETRA products in two *C. reinhardtii* strains (T222+ and CC125+), strongly suggesting that these telomeres have a 3′ overhang of at least 12 nucleotides, corresponding to the size of the annealed part of PETRA-T to the overhang (**Figure 1B** and **Supplemental Figure S1C**). The size of PETRA products (between 800 and 1000 bp) allowed us to evaluate the average length of the 1R, 9R and 10R telomeres by subtracting the distance between the forward primer and the beginning of the telomeric repeats, resulting in telomere lengths of ~700 bp for 1R, ~400 bp for 9R and ~550 bp for 10R in these clones.

As it was shown that a subset of *A. thaliana* telomeres display blunt ends instead of 3′ overhangs (Kazda et al., 2012), we asked whether blunt-ended telomeres also exist in *C. reinhardtii*, since the PETRA experiment alone cannot exclude this possibility. To test this, we applied a hairpin assay, which was successfully used in *A. thaliana* to detect blunt-ended telomeres (Kazda et al., 2012). Briefly, a synthetic hairpin DNA can be ligated to both strands of the telomeres, only if they are blunt-ended. After digestion with *Alu*I at a site in the subtelomeres, the ligated products migrate as a double-sized fragment compared to the unligated control in denaturing conditions. Cleavage of the ligated product by *Bam*HI, using a restriction site designed in the hairpin, can then show that the slow migrating product was indeed generated by ligation to the hairpin (**Supplemental Figure S1D**, left). No blunt ends could be detected using this hairpin assay in two independent biological replicates for strains T222+, CC125+ (**Supplemental Figure 1D**, right), and for four additional strains CC620, CC621, 21gr and 302 (**Supplemental Figure S1E**).

Taken together, the results suggest that most *C. reinhardtii* telomeres end with a 3′ overhang.

### Analysis of *C. reinhardtii* telomeres by <u>T</u>erminal <u>R</u>estriction <u>F</u>ragment (TRF) Southern blots

To study telomere length distributions and their possible regulations, we optimized a TRF Southern blot method for *C. reinhardtii* to accurately measure telomere length from populations of cells. Briefly, a cocktail of six restriction enzymes that do not cut in the canonical and variant telomere motifs of *C. reinhardtii* was selected and predicted to cut ~100 bp from the telomeres, on average. Southern blot analysis of genomic DNA treated by the enzyme cocktail, using a radioactive oligo-probe containing TTTTAGGG telomere repeats allowed for specific detection of telomere-containing fragments (Fulneckova et al., 2013).

We first measured telomere length in three independent biological replicates of strains T222+ and S24-, two isogenic reference strains differing only in their mating-type (Gallaher et al., 2015). We found that telomere fragments spread as a smear over a large range of lengths, from ~200 to ~1200 bp (**Figure 1C**). The two strains displayed a significant difference in their average telomere length (mean ± SD: T222+ = 539 ± 54 bp, N = 18, and S24- = 710 ± 12 bp, N = 5). To demonstrate that the detected smeary signal indeed corresponded to terminal fragments of the chromosomes, we digested the genomic DNA with exonuclease *Bal*31 prior to the digestion with the cocktail of restriction enzymes and Southern blotting (Fajkus et al., 2005; Petracek et al., 1990; Richards and Ausubel, 1988). Briefly, Bal31 degrades both 3′ and 5′ termini of duplexed DNA and thus trims the chromosome extremities without affecting internal regions. We observed that with increasing incubation times with *Bal*31, the signal progressively decreased in size until it nearly disappeared after 10 min (**Figure 1D** and **Supplemental Figure S1F**), demonstrating that it indeed corresponded to terminal telomere sequences. A band at ~200 bp remained unchanged even with the longest Bal31 treatment, indicating that it stemmed from interstitial telomere repeats located within the genome. Since this sharp band did not cross-react with a probe targeting TG microsatellite sequences (**Supplemental Figure S1G**), it most probably corresponded to *bona fide* telomere-sequence-containing region(s) of the genome and not to non-specific cross-hybridizations.

### Telomeres length distribution is stable in different standard growth conditions

*C. reinhardtii* has been widely used as a model organism to study photosynthetic processes due to its ability to grow in different metabolic regimes (Harris, 2009). Under strictly phototrophic conditions (minimum medium in the light), photosynthesis is the only metabolic process providing ATP and reducing power to growing cells. In strictly heterotrophic conditions in the dark, *C. reinhardtii* can survive by respiring the acetate contained in <u>T</u>ris-<u>A</u>cetate <u>P</u>hosphate (TAP) medium. In mixotrophic conditions, *i.e*. TAP medium in the light, cells use a combination of photosynthesis and respiration to grow. Since in other organisms, environmental conditions can regulate telomere length (Epel et al., 2004; Romano et al., 2013; von Zglinicki, 2000; Walmsley and Petes, 1985), we asked whether telomeres vary in length and/or size distributions in response to different standard growth conditions.

We first tested whether cells displayed different telomere lengths during a standard growth kinetic in TAP medium, from inoculation to exponential and then stationary phase, sampled at different time points over a period of 8 days. We observed no significant difference in telomere length between the samples (**Figure 2A**). Prolonged incubation in stationary phase for up to 15 days also did not strongly affect telomere length, despite a slight drop at day 8 (**Figure 2B**). Thus, telomere length were not altered either in exponential growth in replete medium or in the absence of growth, during nutrient depletion and with any other properties of saturated cultures, even over a prolonged period.

**Figure 2:**
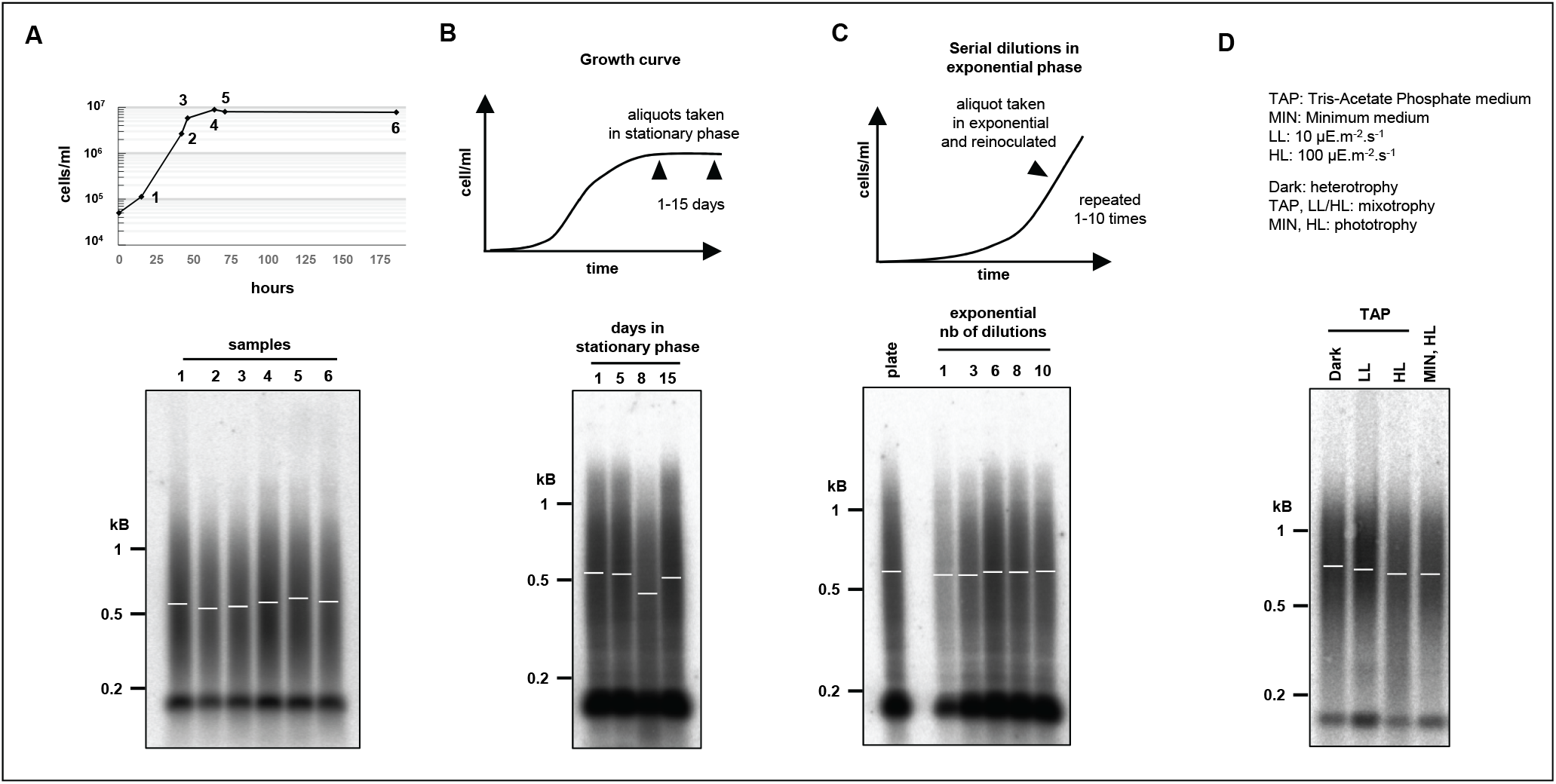
Telomere length distribution is stable under various growth conditions. **(A)** Telomere length distributions of T222+ strain at different growth stages of liquid cultures. T222+ cells were harvested at early exponential (1), mid-exponential (2), late exponential (3) and early (4, 5) and late (6) stationary phases and analyzed by TRF Southern blot. **(B)** Telomere length distributions of prolonged cultures in stationary phase. Cells were harvested after 1, 5, 8 and 15 days after reaching stationary phase. **(C)** Telomere length distributions of serial dilutions of rapidly growing cells. A liquid culture of T222+ cells was grown to exponential phase (2.10^6^ cells/mL), a sample of cells was harvested and the remaining cells diluted with fresh media to 5.10^4^ cells/mL. This serial dilution was repeated 10 times. Samples corresponding to dilutions 1, 3, 6, 8 and 10 were then analyzed by TRF Southern blot. Plate: cells were directly scraped from one week-old streaks on TAP Petri dishes, without liquid culture. **(D)** Telomere length distributions in different metabolic growth conditions. Cells were grown for 6 days to stationary phase either in heterotrophic conditions in TAP medium in the dark, in mixotrophic conditions in TAP medium in low (LL) or higher light (HL), or in pure photo-autotrophic conditions in minimum (MIN) medium under HL.

We also asked whether stimulating cell growth could affect telomere length. Because of the multiple fission mode of cell division of *C. reinhardtii* (Cross and Umen, 2015), actively growing cells might spend less time in each cell cycle and we reasoned that on average telomerase might thus be less active. To test this hypothesis, a TAP culture was constantly maintained in exponential growth phase by serial dilutions over a period of 10 days. Telomere length did not significantly change (**Figure 2C**) and therefore, high division rate did not affect telomere length or distribution.

Finally, we checked telomere length distributions in cultures grown in either strictly phototrophic, strictly heterotrophic, or mixotrophic conditions for 7 days in liquid medium (~20 population doublings) but found no significant difference between the conditions (**Figure 2D**). As telomeres might reach a new steady-state level with a slower kinetic, we repeated the experiment over a period of 60 days (~200 population doublings) but again did not detect changes in telomere length regardless of the growth conditions (**Supplemental Figure S2**).

These experiments demonstrated that *C. reinhardtii* has an active telomere maintenance mechanism and that telomere length distribution is robust with regards to perturbation in metabolic regimes under a variety of standard laboratory growth conditions.

### *C. reinhardtii* reference strains show dramatic differences in telomere length and size distributions

Even though telomere length distribution was very stable under different growth conditions for a given strain (**Figure 2**), we did observe a reproducible and significant difference in mean telomere length between the two laboratory reference strains T222+ and CC125+ by PETRA (**Figure 1B** and **Supplemental Figure S1C**) and between T222+ and S24-by TRF Southern blot (**Figure 1C**). We thus wondered if closely related but divergent *C. reinhardtii* strains displayed significant inter-strain differences in telomere length distributions. To test this, we took advantage of the recent sequencing of many closely related reference strains widely used in different laboratories across the world and which display up to 2% genetic divergence (Gallaher et al., 2015). We performed TRF Southern blots on 12 related but divergent *C. reinhardtii* strains to characterize their telomeres (**Figure 3A**). Strikingly, steady-state telomere lengths were highly variable from strain to strain, ranging from 378 ± 24 bp (mean ± SD, N = 4) in CC125+ to 3.2 ± 1.1 kb (N = 3) in cw15.J14+, encompassing nearly one order of magnitude (**Figure 3B**). Telomere length did not correlate with genome divergence (genetically close strains are depicted with the same color) and we did not find any obvious genomic region, as described in Gallaher *et al*. (2015), that would co-segregate with longer or shorter telomeres. In particular, neither the mating type, nor the presence or absence of a cell wall correlated with telomere length variations. The average telomere length in strain cw15.J14+ was particularly striking and we asked whether the signal corresponded to internal telomere repeats. A Bal31 exonuclease treatment time course prior to TRF Southern blotting showed the signal decreasing in size demonstrating that this signal indeed corresponded to terminal repeats (**Supplemental Figure S3B**, right). In addition to length variations, some strains, such as CC503+ and CC1010+, displayed multimodal telomere length distributions (**Figure 3A, 3B** and **Supplemental Figure S3A**), some peaks of which might correspond to internal telomere repeats. To test this possibility, we performed a *Bal*31 treatment experiment on strain CC503+, prior to TRF analysis. The whole smear, including the three peaks of the multimodal distribution, was progressively degraded with increasing digestion time, demonstrating that the multimodal distribution corresponded to terminal telomere repeats of different lengths (**Supplemental Figure S3B**, left).

**Figure 3:**
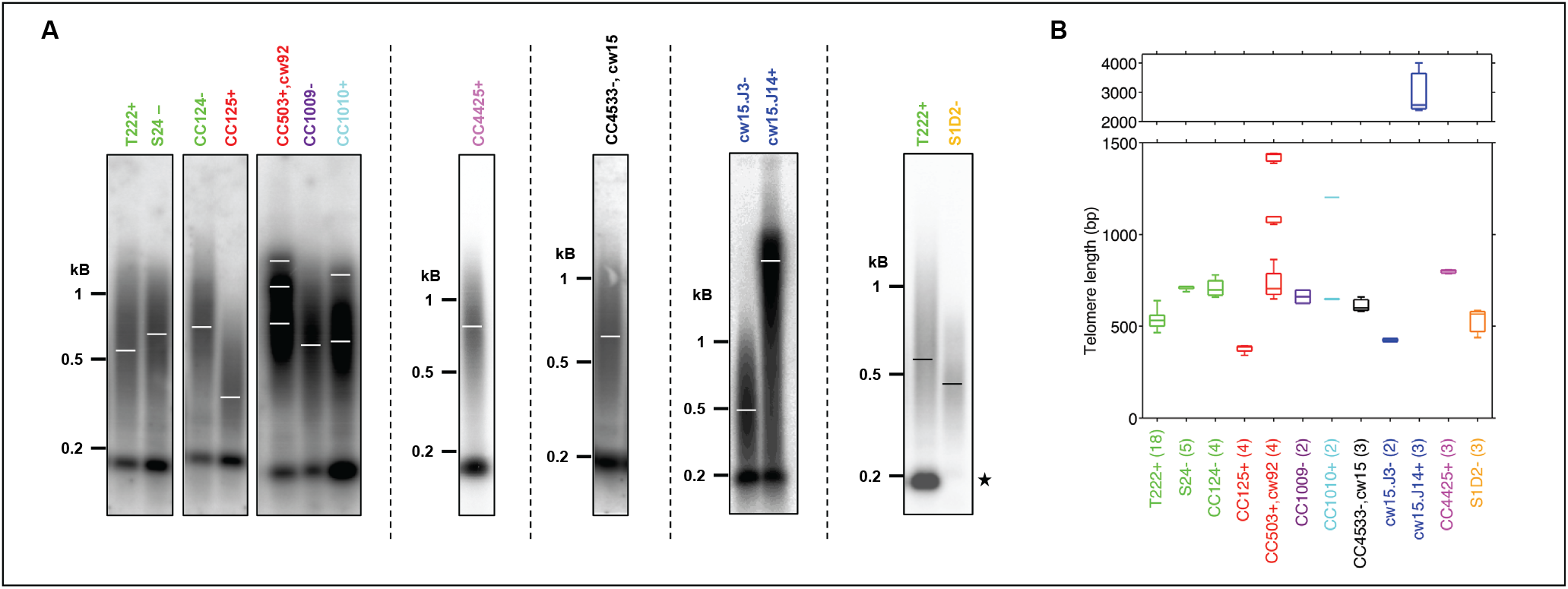
Vast differences in telomere length distributions in *C. reinhardtii* reference strains. **(A)** Telomeres of recently sequenced *C. reinhardtii* reference strains (Gallaher *et al*., 2015) were analyzed by TRF Southern blot. Strains sharing the same name color are closely related genetically, while strains with different colors are more divergent. Dashed vertical lines indicate independent gels. Star: S1D2-strain does not display the band at ~200 bp. cw15 and cw92 indicate mutations that led to cell-wall-less strains. **(B)** Mean and standard deviation of telomere length for each strain as calculated by analysis of Southern blots from the indicated number of independent biological replicates biological replicates (N).

Interestingly, the interstitial band at ~200 bp, which was present in 11 tested *C. reinhardtii* reference strains was absent from the S1D2- (CC2290-) strain. S1D2- is an interfertile but divergent *C. reinhardtii* species, often used for genetic mapping purposes (Gross et al., 1988; Vysotskaia et al., 2001). Thus, the interstitial telomere sequence might have emerged in a subset of *C. reinhardtii* species or conversely might have been lost in S1D2-.

### Identification of the gene encoding the catalytic subunit of telomerase

Telomerase is a holoenzyme comprised of at least a reverse-transcriptase catalytic subunit and a template RNA, which are sufficient for *in vitro* telomerase activity (Lingner et al., 1997a). These core actors are associated with multiple other proteins, required for its recruitment, processivity and regulation (Lewis and Wuttke, 2012). As the catalytic subunit of telomerase (*e.g*. hTERT in human, AtTERT in *A. thaliana* and Est2 in *S. cerevisiae*) is conserved, we sought to identify the gene encoding this subunit in *C. reinhardtii* and to characterize the contribution of telomerase to telomere length maintenance.

Gene model Cre04.g213652 of the *C. reinhardtii* nuclear genome (Phytozome v5.5; https://phytozome.jgi.doe.gov/pz/#) has a predicted N-terminal part of the corresponding protein showing partial sequence similarity with RNA-binding domains of telomerase from a number of organisms (**Figure 4A**). The available gene model extends over 25 kb, contains 28 introns and is predicted to encode a 5019-aa protein, much larger than telomerases from *A. thaliana* (1123 aa), maize (1188 aa), iris (1295 aa) and rice (1261 aa). Two sequencing gaps and the presence of TG and CCAC satellites in the gene model (both in introns and in exons) cloud the structure of the putative gene. While expressed sequence tags from cDNA libraries supported the validity of some parts of the conserved 5′ and 3′ regions, no expressed sequence tag was found for the central part of the gene model in the available *C. reinhardtii* expression libraries. Nucleotide sequence alignments failed to detect similarity with telomerase catalytic subunit genes of other organisms. We thus performed PSI-Blast alignments of the C-terminal protein domain of the putative *C. reinhardtii* telomerase with telomerases from plants using PRALINE (http://www.ibi.vu.nl/programs/pralinewww). The alignments showed strong similarity to the C-terminal catalytic reverse transcriptase domain of *A. thaliana* (e-value = 3.10^−36^), maize (e-value = 4.10^−35^), iris (e-value = 1.10^−36^) and rice (e-value = 3.10^−24^) (**Figure 4B**). The conserved C motif (mC) in telomerases ranging from *S. cerevisiae* to *A. thaliana* and humans including the two critical aspartates for telomerase catalytic activity (Lingner et al., 1997b; Nakamura et al., 1997; Oguchi et al., 1999) showed strong sequence conservation with a corresponding motif in the putative *C. reinhardtii* protein (**Figure 4B** and **4C**). Motif E (mE) was conserved to a lesser degree, while no clear conservation of motifs mA, mD, motif 1 and 2 (Lingner et al., 1997b; Oguchi et al., 1999) was found in the predicted *C. reinhardtii* protein. Other well conserved regions in the C-terminal part with no assigned motif are also depicted in **Figure 4B**.

**Figure 4:**
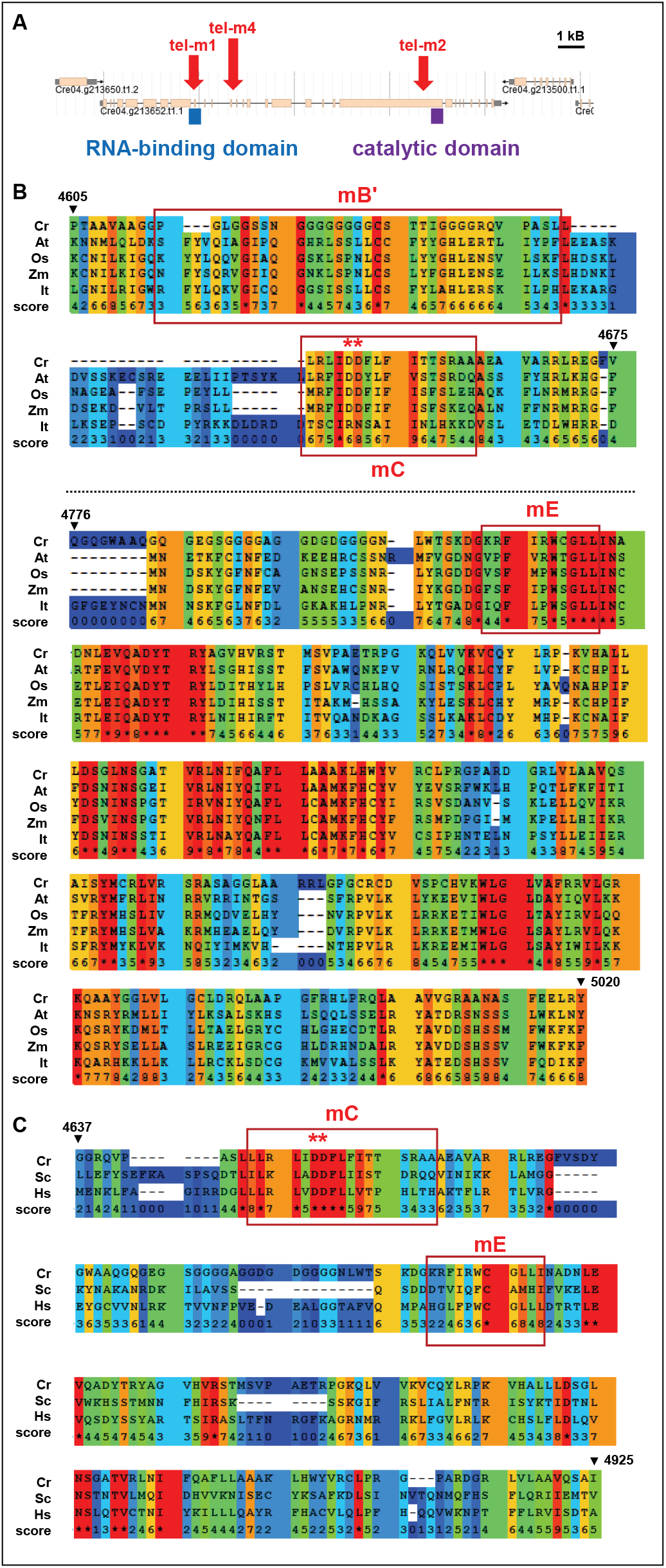
Identification of the *CrTERT* gene encoding the catalytic subunit of telomerase in *C. reinhardtii*. **(A)** The protein corresponding to the predicted gene model Cre04.g213652.t1.1 of the available *C. reinhardtii* nuclear genome harbors an annotated N-terminal domain with significant similarities to the RNA template-binding domain of telomerases from other organisms. The C-terminal domain shows strong similarities with the catalytic domain of this enzyme in other organisms. Mutants tel-m1 (LMJ.RY0402.077111) and tel-m2 (LMJ.RY0402.209904) from the CliP library have reported insertions in either the RNA-binding or the catalytic domain, respectively. Mutant tel-m4 (LMJ.RY0402.105594) has an insertion in between these two domains. **(B)** PSI-blast alignments show strong amino-acid sequence similarity of the catalytic domain of telomerases from many organisms with the putative *C. reinhardtii* protein. Similarity score ranges from 0 (light blue) to 9 and * (red) indicates identity. *Cr, C. reinhardtii; At, A. thaliana; Os, O*. sativa; *Zm, Z. mays; It, I. tectorum*. The motifs B’, C and E (mB’, mC and mE) described in (Lingner et al. 1997, Ogushi *et al*. 1999) show strong conservation in *C. reinhardtii*, including two catalytic aspartates, essential for telomerase function in other organisms (red star). Conservation can also be observed downstream of mE between *CrTERT* and the other telomerases. **(C)** The mC motif of *C. reinhardtii* shows strong sequence similarity with the mC motif containing two catalytically essential aspartates in yeast and human telomerases (Lingner et al. 1997, Ogushi *et al*. 1999).

To demonstrate that the genomic region Cre04.g213652 indeed contains the gene encoding the catalytic subunit of telomerase of *C. reinhardtii*, we used a genetic approach. We selected three strains harboring insertions of the paromomycin resistance cassette within the putative gene from the recently created CliP library of mapped insertional mutants (Li et al., 2016) (https://www.chlamylibrary.org) (**Figure 4A**). LMJ.RY0402.077111 has an insertion in a putative intron near the region encoding the putative RNA-binding domain of the gene and was named tel-m1. LMJ.RY0402.209904 has an insertion in the putative CDS of the putative catalytic C-terminal domain and was named tel-m2. LMJ.RY0402.105594 has an insertion in an intron in a non-conserved region between these two domains and was named tel-m4. Although the insertions in these three mutants were already mapped by the work of Li *et al*. (2016) with a confidence of 95% for tel-m1 and tel-m4 and 73% for tel-m2, we verified that all three mutants indeed had the insertion at the predicted loci, using PCR with primers targeting the gene and/or the inserted paromomycin resistance marker (**Supplemental Figure S4A** and **S4B**). For all three mutants, the obtained PCR products were gel-excised, sequenced and shown to correspond to the expected genomic region. We also backcrossed mutants tel-m1 and tel-m2 with the paromomycin-sensitive T222+ strain and analyzed the segregation of the paromomycin resistance phenotype in tetrads after sporulation of the diploids. Correct 2:2 segregation of the mating locus in the offspring of the tetrads was checked by PCR (**Supplemental Figure S4D**). Paromomycin resistance systematically segregated with a 2:2 ratio in the haploid offspring, suggesting that the functional marker was not inserted at multiple loci in the genome (**Supplemental Figure S4C**).

We then analyzed the telomere length of the three mutant strains. All three mutants showed significantly shorter telomeres when compared to the parental CC4533-strain used by Li *et al*. (2016) to construct the CliP library (**Figure 5A** and **Supplemental Figure S5A**; mean ± SD, tel-m1: 373 ± 25 bp, N = 4, tel-m2: 383 ± 30 bp, N = 4, and tel-m4: 387 ± 12 bp, N = 2, compared to CC4533-: 614 ± 41 bp, N = 3). We verified that the shorter telomere length in mutants tel-m1, tel-m2 and tel-m4 was not simply due to the transformation protocol used to generate the CliP library or to the insertion of the paromomycin marker itself. Telomere length was measured in another mutant from the CliP library, harboring an insertion elsewhere in the genome (on chromosome 1), and was comparable to the parental CC4533-strain (**Figure 5A**, “control” and **Supplemental Figure S5D**).

**Figure 5:**
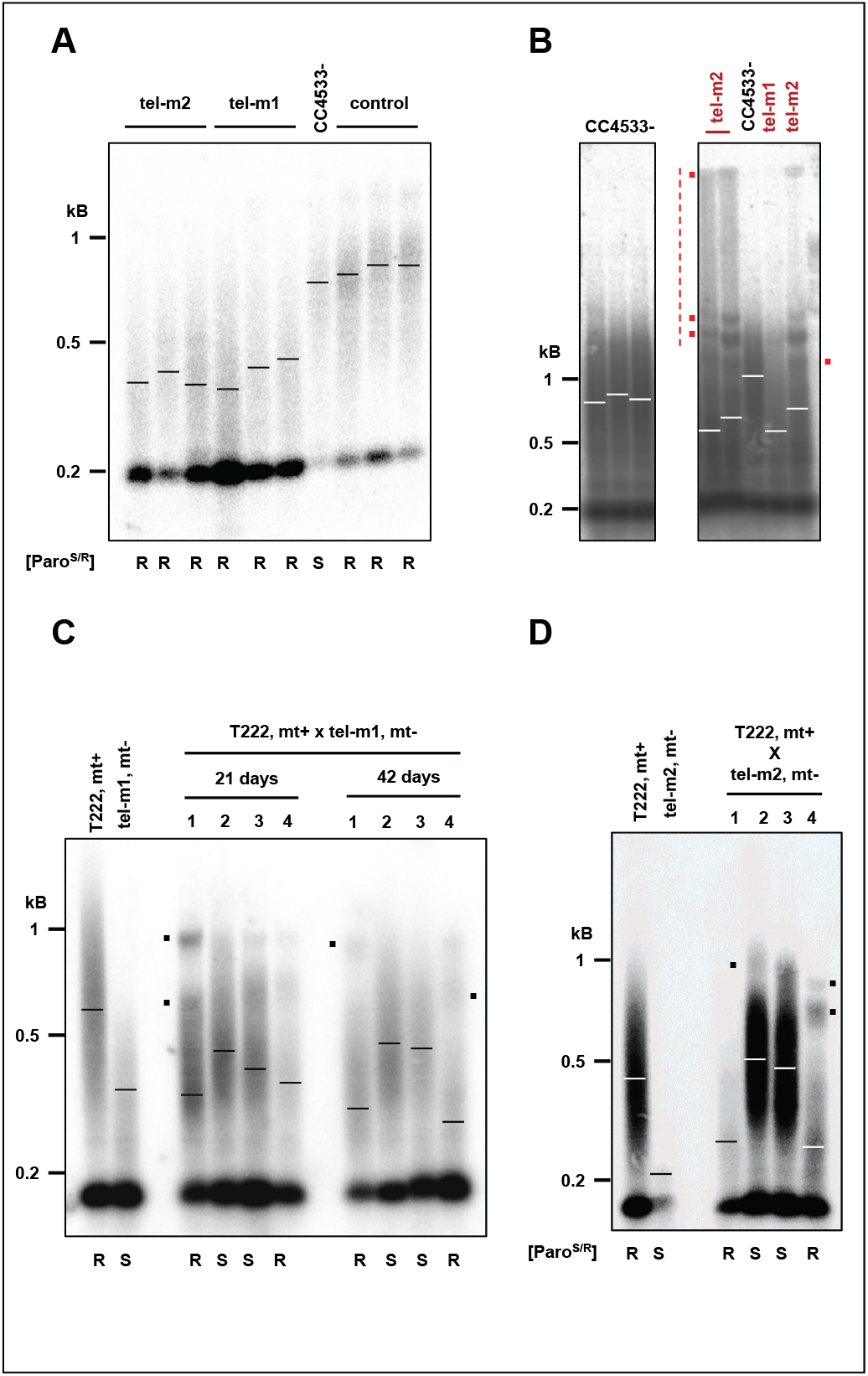
Insertional mutants of the *CrTERT* gene have shorter telomeres. **(A)** Mutants tel-m1 and tel-m2 have shorter telomeres in TRF analyses (three independent subclones are shown). Control: mutant LMJ.RY0402.239308 from the CliP library, which has an insertion in a gene unrelated to *CrTERT*. Paromomycin resistance phenotype is indicated (“[Paro^S/R^]”; “S”: sensitive, “R”: resistant). **(B)** Prolonged liquid cultures of telomerase mutants lead to rearranged TRF patterns. Cells were cultured in liquid medium for two months before TRF analysis. Additional bands and slow migrating DNA molecules (red dots and dotted vertical line, respectively) are indicated for tel-m1 and tel-m2, and are not present in the CC4533-reference strain TRF pattern. **(C)** Tetrad analysis of the cross between tel-m1 and T222+ shows a 2:2 co-segregation of paromomycin resistance and shortened telomeres after 21 and 42 days after the cross (see also **Supplemental Figure S5B**). **(D)** Tetrad analysis of the cross between tel-m2 and T222+ shows a 2:2 co-segregation of paromomycin resistance and shortened telomeres after ~80 days after the cross (see also **Supplemental Figure S5C**).

We conclude that while gene model Cre04.g213652 might be wrong in its predicted structure and will require further study to be corrected, this genomic region indeed harbors the gene encoding for the catalytic subunit of telomerase in *C. reinhardtii*, and we propose to rename the gene model *CrTERT*.

### Telomere rearrangement and maintenance in long-term cultures of telomerase mutants

Since telomeres shortened in telomerase-negative cells, we wondered whether the cells would experience replicative senescence after an extended period of growth, when telomeres reach a critically short length. We thus grew the telomerase mutants tel-m1 and tel-m2 as well as the reference strain for two months (~200 population doublings), with dilutions into fresh TAP medium every 5 days. While we did not observe *CrTERT* mutant cultures dying out and no obvious growth defect was detected at any time point, TRF analysis of the telomere length distribution of tel-m1, tel-m2 showed a drastic change in telomere length distribution (**Figure 5B**, compare with **Figure 5A**): first, the bulk of the telomeres seemed to have a short average length but longer than initially (~500 bp, compared to mean ± SD = 383 ± 30 bp); secondly, additional discreet bands appeared at sizes above 1 kb (red dots); finally, a signal that extended up to the wells was detected (vertical red line). Interestingly, the three independent cultures of the tel-m2 mutant gave similar but distinct patterns with respect to the discreet bands and the high molecular weight signal. The tel-m1 mutant also showed on average longer telomeres after an extended period of culture than initially (**Figure 5B**, compare with **Figure 5A**) and displayed some additional bands, albeit not to the extent of tel-m2. Overall, these altered TRF patterns observed in prolonged cultures of telomerase mutants are reminiscent of TRF patterns observed for cells with telomerase-independent maintenance pathways (*e.g*. type II survivors of telomerase-negative yeast cells or ALT-like telomerase-negative cancer cells. See discussion.)

### Telomeres shorten progressively in telomerase mutants

The initial CliP telomerase mutants might have accumulated additional, potentially suppressor, mutations, which could interfere with the proper assessment of the mutant phenotype. Importantly, the presence of suppressive mutations could explain why these mutants did not show any discernable growth defects in standard growth conditions or any sign of senescence after prolonged culture.

To outcross potential suppressor mutations and gain a kinetic perspective on telomere shortening in the telomerase mutants, we backcrossed mutants tel-m1 and tel-m2 with a wild-type strain of opposite mating type (T222+) and, after sporulation of the diploids, studied the telomere length distribution of the obtained tetrads. Backcrossing a mutant cell with a telomerase-positive strain should allow telomerase to elongate the shortest telomeres brought in by the mutant strain. The subsequent meiosis would then shuffle the chromosomes and the telomeres in the spores, independently of the mutant or wild-type status of the telomerase gene. We thus expect that immediately after sporulation of the diploid, the four spores would have similar and nearly wild-type average telomere length. After culture, the telomere length in the four progenies should vary according to the status of the *CrTERT* gene.

Strikingly, measurement of telomere length in the four haploid progenies of the tel-m1- x T222+ cross after 21 days showed that two of them displayed longer average telomere length and the other two shorter telomeres, which corresponded to the telomerase mutants as assessed by paromomycin resistance (**Figure 5C**, “21 days”). After 21 more days, the telomeres of the telomerase-positive cultures maintained their average length, whereas the telomerase-negative cultures displayed further shortening of their telomeres (**Figure 5C**, “42 days”, and **Supplemental Figure S5B**). Therefore, mutation of the *CrTERT* gene led to an “Ever Shorter Telomere” (EST) phenotype as first described in *S. cerevisiae* (Lundblad and Szostak, 1989). A similar result was observed for the progenies of the cross tel-m2- x T222+ (**Figure 5D** and **Supplemental Figure S5C**). These results strongly argued against the possibility that the shorter telomeres observed in tel-m1 and tel-m2 were due to additional mutations in the genome, because they would not necessarily have co-segregated with the paromomycin marker. We also noted the presence of other bands and peaks in the smear, which were likely the result of segregating parental telomeres of very different lengths during meiosis (black dots in **Figure 5C, 5D** and **Supplemental Figure S5B**).

While no growth defects were observed for the initial tel-m1 and tel-m2 mutants, analysis of the progeny of the spores from backcrosses between tel-m1 and tel-m2 with the wild-type T222+ strain (n = 4 independent tetrads, with 8 telomerase-negative spores) showed that 4 out of the 8 telomerase-negative haploid progenies experienced growth defects and then massive cell death, typical of replicative senescence (highlighted in red in the table of **Supplemental Figure S5E**). Strikingly, for each of these 4 telomerase-negative progenies that experienced massive cell death, some cells managed to form colonies again at very low frequency (**Supplemental Figure S5E**, left) and thus corresponded to post-senescence survivors. The 4 other telomerase-negative haploid progenies did not display any growth defect (highlighted in green in the table of **Supplemental Figure S5E**). Individual colonies of post-senescent survivors kept on solid media showed cycles of moderate growth and subsequent cell death. This complex and dynamic survivor phenotype will be investigated in future studies.

## DISCUSSION

In this study, we provide a detailed molecular characterization of *C. reinhardtii* telomeres by investigating their sequence, end-structure and length distribution. We also identify *CrTERT*, the gene encoding the catalytic subunit of telomerase, and find that mutants of this gene display an “Ever Shortening Telomere” phenotype and can enter replicative senescence.

### Telomere repeats and variants

A precise knowledge of the sequence architecture of telomeric repeats in *C. reinhardtii* is important information for the understanding of the molecular mechanisms underlying their physiological roles (*e.g*. shortening, lengthening, gene expression regulation and binding of regulatory proteins). Some species such as *S. cerevisiae* and *S. pombe* display degenerated telomere sequences, while other organisms harbor mostly identical repeats (Zakian, 1995). Because telomere-bound proteins specifically interact with telomere sequences (Fulcher et al., 2014; Palm and de Lange, 2008), the variability of telomeric repeat motif can have functional consequences. For example, the presence of sequence variants can create binding sites for other proteins: in *S. cerevisiae*, the presence of the human type TTAGGG motif close to the (TG_1-3_)_n_ telomeres creates binding sites for the essential Tbf1 transcription factor, which contributes to telomerase recruitment and may provide an alternative capping (Arneric and Lingner, 2007); and in human cancer cells with alternative telomere maintenance mechanism, variant telomere sequences are bound and inserted in the genome by nuclear receptors, which destabilizes the genome (Marzec et al., 2015).

By analyzing 32 independent clones and 709 telomeric repeats, we come to the conclusion that *C. reinhardtii* telomeric repeats are mostly non-degenerated, with few low-frequency variants, notably repeats of the canonical *A. thaliana* type (TTTAGGG). This repeat is also found as the first repeat in 10 subtelomere-telomere junctions from eight chromosomes (**Table 1** and **Supplemental Figure S1A**), possibly a remnant of the ancestral motif in the green lineage (Fulneckova et al., 2012). The low occurrence of other variants (**Table 1**) suggests that *C. reinhardtii* telomerase is a high fidelity reverse transcriptase, in contrast to telomerase from other unicellular eukaryotes such as *S. pombe* or *S. cerevisiae*.

Analysis of the available genome sequences of *C. reinhardtii* strains shows some occurrences of interstitial telomeric repeats and our TRF Southern blot experiments showed a non-terminal fragment of ~200 bp, the length of which is defined by its resistance to digestion with the cocktail of restriction enzymes we used (**Figure 1D** and **Supplemental Figures S1F and S3B**, denoted by a star). The presence of the interstitial telomeric repeats might be due to chromosome end-to-end fusion over the course of evolution (Aksenova et al., 2015; Azzalin et al., 2001; Gaspin et al., 2010; Meyne et al., 1990; Uchida et al., 2002). Furthermore, as telomere sequences are binding sites for specific proteins, which may act as transcription factors (*e.g*. Rap1 in yeast), they can act as transcriptional regulators of intragenomic loci (Platt et al., 2013).

### Intra-strain stability and dramatic inter-strain variations in telomere length distributions

Telomere length is regulated by multiple pathways, as shown by exhaustive screens performed in *S. cerevisiae* (Askree et al., 2004; Chang et al., 2011; Gatbonton et al., 2006; Ungar et al., 2009). These pathways are very diverse and include nucleic acid metabolism, DNA replication, chromatin modification and protein degradation, among others. In addition, telomere length is also sensitive to both internal and environmental cues (Cetin and Cleveland, 2010; Epel et al., 2004; Fulcher et al., 2014; Millet et al., 2015; Millet and Makovets, 2016; Romano et al., 2013; von Zglinicki, 2000; Walmsley and Petes, 1985). We found no change in telomere length distribution when *C. reinhardtii* cells were grown in a wide variety of standard laboratory conditions, including growth phases (exponential vs stationary), carbon source and light conditions, which are all relevant for physiological growth of this alga. While we cannot exclude that other harsher growth conditions or internal signaling (*e.g*. DNA damage or replication stress) might induce an alteration in telomere length or structure, this result suggests that the mechanisms maintaining telomere length homeostasis are highly robust and efficient.

In stark contrast, closely related strains of *C. reinhardtii* displayed very different telomere length profiles, suggesting that steady-state telomere length is not under selective pressure. In particular, strains with very short (*e.g*. CC125+) or very long (*e.g*. cw15.J14+) telomeres (**Figure 3**) might differ in their telomerase activity or in other regulators of telomere homeostasis. However, no obvious genome region could be correlated to this length variation. A more detailed functional genetic approach to map the regions of the genome responsible for telomere length variation could identify pathways regulating telomere length.

Beside length variation, some strains such as CC503+ and cw15.J14+ displayed multimodal profiles (**Figure 3**). Such profiles could be explained by a heterogeneous cell population with a subpopulation of cells harboring very different average telomere lengths. Given that all our experiments were performed after subcloning, the phenotypic heterogeneity could have arisen in a genetically uniform population of cells and been maintained as an epigenetic trait. Another hypothesis would be that different telomeres within a cell might have different steady-state lengths, possibly through local cis-regulation mechanisms.

Overall, the diversity of telomere length distributions observed in these reference strains highlights the plasticity of telomere length regulation and the phenotypic heterogeneity of *C. reinhardtii* reference strains.

### Identification of *CrTERT* encoding the catalytic subunit of telomerase

Based on sequence similarity (**Figure 4**) and functional analyses of three independent mutant alleles of the gene Cre04.g213652 (**Figure 5** and **Supplemental Figure S5**), we propose that it corresponds to, or at least encompasses, the gene encoding the catalytic subunit of telomerase, required to maintain telomere length in *C. reinhardtii*. We propose to rename it *CrTERT*.

Multiple lines of evidence support this conclusion. First, the predicted protein shares significant sequence similarity with the RNA-binding domain of telomerase from other organisms in its N-terminus. Secondly, we find a very strong conservation of the C-terminal domain of the proposed CrTERT protein with catalytic domains of telomerases not only from plants (Maize, Arabidopsis, Soya, Iris; **Figure 4B**) but also from yeast and human (**Figure 4C**). Motif C (mC) is particularly well conserved, including two aspartates essential for the catalytic activity of telomerase (**Figure 4B** and **4C**). Thirdly, three independent mutants (tel-m1, tel-m2 and tel-m4) bearing different insertions of the paromomycin resistance marker in *CrTERT*, including within its RNA-binding domain (tel-m1) and its catalytic domain (tel-m2) display significantly shorter telomeres than the parental CC4533-strain, which is not the case for other independent mutants from the CliP library located in loci unrelated to telomerase (**Figure 5A, Supplemental Figure S5D**). Backcrosses of tel-m1 and tel-m2 showed a 2:2 segregation of paromomycin resistance associated with shorter telomere lengths (**Figure 5C** and **5D**; **Supplemental Figure S5B** and **S5C**), indicating that a single insertion of the paromomycin marker in *CrTERT* was responsible for the observed phenotype. Finally, telomeres shortened progressively in paromomycin-resistant progenies. However, as we are as of yet unable to detect the mRNA corresponding to *CrTERT* by either northern blotting or RT-qPCR, possibly because of its low expression, we could not assess *CrTERT* expression in our study.

The identification of additional components of the telomerase holoenzyme and telomere associated proteins will be the focus of future work.

### Telomere shortening, replicative senescence and alternative maintenance pathways

After prolonged liquid cultures of multiple independent tel-m1 and tel-m2 mutants, we observed a drastically altered TRF pattern: discrete bands above the 1.5 kb range (**Figure 5B** red dots) as well as a continuous smear of high molecular weight fragments up to the wells (**Figure 5B**, vertical red line). These new TRF signals could correspond to extremely long telomeres, as seen for strain cw15.J14+, but also to DNA molecules with abnormal structures, such as G-quartets, other secondary structures or single-stranded DNA. These rearrangements suggest that alternative mechanisms of telomere maintenance or elongation might have been activated or selected. Overall, the altered telomere length distribution in long-term cultures of *CrTERT* mutants is reminiscent of telomere profiles observed in type II post-senescent yeast cells (Lundblad and Blackburn, 1993), ALT (Alternative Lengthening of Telomeres) cancer cells (Cesare and Reddel, 2010; Shay et al., 2012) or ALT *A. thaliana* cell lines (Akimcheva et al., 2008; Zellinger et al., 2007), in which telomerase-independent recombination mechanisms can lead to very long and heterogeneous telomeres, thus sustaining long-term cell divisions. In these described cases, telomerase is not expressed, telomeres undergo sister-chromatid and inter-chromosome homologous recombination using gene conversion, break-induced replication, rolling circle amplification or yet unknown mechanisms. This telomerase-independent telomere elongation leads to a change of telomere and subtelomere structures, revealed by distinct TRF patterns, resembling the ones we observe after extended culture of the tel-m1 and tel-m2 mutants.

Another line of evidence suggesting the occurrence of post-senescence survivors of telomerase-negative cells in *C. reinhardtii* came from the analysis of the offspring of backcrosses of the tel-m1 and tel-m2 mutants with T222+ reference strain. The *CrTERT* mutant spore progenies displayed an “Ever Shortening Telomere” phenotype (**Figure 5C** and **5D**; **Supplemental Figure S5B** and **S5C**) and 50% of them eventually stopped growing after about 6 months on solid media, a phenotype consistent with replicative senescence (**Supplemental Figure S5E**). The other 50% of telomerase-negative spores has not entered senescence as of yet (> 18 months). The spores that experienced senescence and generated first generation survivors then showed a complex pattern of moderate growth, followed by cell death and emergence of a new generation of clonal survivors. We do not yet understand the variability of the senescence phenotype in these backcrossed haploid progenies. We speculate that for the initial CliP mutants, additional mutations could have been generated that might have acted as suppressors of the senescence phenotype. This would also explain why no growth defects were observed for the initial CliP mutants even after more than two years of maintenance on solid media, while senescence, cell death and post-senescent survivors could be observed after backcrossing these mutants and selecting telomerase-negative spore progenies. Alternatively, the initial CliP mutants might have already been post-senescence survivors from the beginning. In a future work, it will be interesting to characterize post-senescence survivors by assessing hallmarks of human ALT cancers, including for example circular extrachromosomal telomeric DNA and up regulation of telomeric repeat-containing RNA (TERRA) (Arora and Azzalin, 2015; Cesare and Reddel, 2010).

While some fundamental aspects of its telomeres share similarities to other eukaryotes, *C. reinhardtii* shows a unique combination of telomeric properties that distinguishes it from any other model organism. The characterization of its telomeres at the level of sequence, end-structure, length distribution and maintenance by telomerase or alternative mechanisms, provided by this study is an essential step to propose *C. reinhardtii* as a valuable model organism for telomere biology research.

## METHODS

A detailed description of the methods used can be found in Supplemental Methods.

## AUTHOR CONTRIBUTIONS

Conceptualization: SE, SV, FAW, MTT, KR and ZX. Supervision: SE, KR and ZX. Investigation: SE, SV, JR, PJ, SB and ZX. Formal Analysis: all authors. Writing – Original Draft: SE, SDL and ZX. Writing – Review & Editing: all authors.

## ACKNOWLEDGMENTS

We thank the Chlamydomonas Mutant Library Group at Princeton University, the Carnegie Institution for Science, and the Chlamydomonas Resource Center at the University of Minnesota for providing the indexed Chlamydomonas insertional mutants. This work was supported by the ANR grant “AlgaTelo” (ANR-17-CE20-0002-01) to ZX, la Fondation de la Recherche Médicale (MTT “équipe labellisée”), the ANR grant “InTelo” (ANR-16-CE12-0026) to MTT, the ‘‘Initiative d’Excellence’’ program from the French State (Grant ‘DYNAMO’, ANR-11-LABX-0011) and by the Ministry of Education, Youth and Sports of the Czech Republic, European Regional Development Fund-Project “REMAP” (No. CZ.02.1.01/0.0/0.0/15_003/0000479) to KR. The authors declare no conflict of interest.

