## Supplemental Information for "Molecular characterization of *Chlamydomonas reinhardtii* telomeres and telomerase mutants"

**Supplemental Figure S1: Characterization of *C. reinhardtii* telomere repeats by telomere-PCR and PETRA. Related to Figure 1. (A)** Scheme of the 17 *C. reinhardtii* chromosomes. Red arrow: forward primer oT1090 used in telomere-PCR. Blue arrows: forward primers 1R, 9R and 10R used in PETRA experiments. Green numbers refer to telomere length from the public genome sequence of *C. reinhardtii* (Phytozome genome version 5.5; https://phytozome.jgi.doe.gov/pz/#!info?alias=Org_Creinhardtii). ND: no telomere in the database. **(B)** Scheme of the different steps of telomere-PCR (left). After adding a poly-C extension to telomeric ends by terminal transferase, PCR amplification using a forward primer at a subtelomere-telomere junction (oT1090) and a poly-G reverse primer was performed. The resulting PCR products were cloned and sequenced. Telomere-PCR was used to amplify telomeric repeats from the T222+ strain. The resulting PCR products were cloned in *E. coli* and 32 clones sequenced. Five examples of telomere-PCR sequences showing the sequence of primer oT1090 (yellow), the poly-C extension (blue), canonical *C. reinhardtii* repeats (TTTTAGGG, in red) and variant motifs (other colors) (right). The frequency of the observed repeats is indicated in **Table 1**. **(C)** Scheme of the PETRA assay for 3′ overhang detection. The primer PETRA-T was annealed to the putative 3′ overhang and extended using DNA PolI. PCR amplification was then performed using a subtelomere-specific primer and PETRA-A primer, which annealed to the non-telomeric sequence of the 5′ end of PETRA-T. (top). An independent biological replicate of the result shown in **Figure 1B** is shown at the bottom. **(D)** Left: scheme of the hairpin assay to detect blunt-end telomeres as described (Kazda et al., 2012). Right: two independent biological replicates for strains T222+ and CC125+ show no evidence for blunt telomeres. **(E)** Four other *C. reinhardtii* strains CC-620, CC-621, 21gr and 302, were tested and were negative for blunt ends by hairpin assay. **(F)** Independent biological replicate of the *Bal*31 assay performed on strain T222+ (see **Figure 1D**). **(G)** A poly-TG probe did not detect the band observed at ~200 bp with a telomere probe. Genomic DNAs from strains T222+, S24-, CC124- and CC125+ were digested with the same protocol used for TRF experiments, Southern blotted and then probed with a radioactive $(TG)_{13}$ probe. T222+ samples were also treated by *Bal*31 exonuclease for increasing amounts of time (5 first lanes) before digestion with the restriction enzymes. NP1-3 samples were not column-purified.

**Supplemental Figure S2: Prolonged culture in different growth conditions does not significantly affect telomere lengths. Related to Figure 2.** Reference strains T222+ and S24- were grown in either heterotrophic conditions (TAP, Dark) or in photo-autotrophic conditions (MIN, HL). When reaching the stationary phase, cultures were diluted in fresh media and this was repeated for 2 months, *i.e.* ~200 population doublings (PD) before TRF analysis. Triangle: band not seen in other TRF experiments.

**Supplemental Figure S3: Multimodal and very long telomeres from CC503+ and cw15.J14+ correspond to terminal DNA fragments. Related to Figure 3. (A)** TRF Southern blot of biological replicates of the indicated strains shown in **Figure 3** (left), with intensity profile analysis (right) showing the multimodal distribution of CC503+ and CC1010+. **(B)** Genomic DNAs of strains CC503+ and cw15.J14+ were subjected to *Bal*31 exonuclease digestion for 1 to 10 minutes prior to digestion by the restriction enzyme cocktail and Southern blotting using the CHSB (oT0958) probe, showing that the detected signal is indeed *Bal*31-sensitive and hence represents terminal fragments.

**Supplemental Figure S4: PCR verification of insertion mutants tel-m1 and tel-m2. Related to Figure 4**. **(A)** Scheme of expected PCR amplification product (left). $F_x$ and $R_x$ denote primers where "x" stands for "m1" or "m2". PCR using primers specific to the reported insertions sites in tel-m1 ($F_{m1}$ and $R_{m1}$) and tel-m2 ($F_{m2}$ and $R_{m2}$) gave no signal in the targeted mutant because the inserted marked is loo large for efficient amplification, but amplified the expected product in the reference strain and the other mutant (of size 1.8 kb and 571 bp, respectively; gels in the middle and on the right). Primers $F_c$ and $R_c$ amplified of a 422-bp product in the *CrTERT* gene outside of the predicted insertion sites in tel-m1 and tel-m2 (positive control, gel on the left). **(B)** Scheme of expected PCR amplification product using specific primers for tel-m1 ($F_{m1}$) and tel-m2 ($F_{12}$), combined with primers in the paromomycin cassette ($K_1$ and $K_2$) (left). The two pairs of primers amplified the expected products in the respective mutants (right). All bands were excised, gel-purified and sequenced**.** PCR-amplification using primers $K_1$ and $R_{m4}$ were used to characterize the insertion in tel-m4. The specific band obtained in tel-m4 was excised from the gel and sequenced and could indeed be mapped to the expected *CrTERT* sequence. **(C)** Mutant tel-m1 was crossed with T222+ strain. The cross with tel-m2 is not shown. The parental strains (tel-m1 and T222+) as well as the progeny of the four spores of a tetrad from the diploid are spotted on TAP solid media with ("+ Paromomycin") or without paromomycin (10 μg.mL$^{-1}$). Five tetrads were analyzed for each cross for the segregation of paromomycin resistance. **(D)** The 2:2 segregation of the mating type was checked by PCR.

**Supplemental Figure S5: Telomere length and senescence phenotype of *CrTERT* mutants. Related to Figure 5. (A)** Insertion mutant tel-m4 has shorter telomeres than T222+ and CC4533- as shown by TRF Southern blots of two biological replicates of the tel-m4 mutant strain. **(B)** Independent tetrad of the tel-m1 x T222 cross analyzed by TRF Southern performed on the spore progenies (see also **Figure 5B**). **(C)** Independent tetrad of the tel-m2 x T222 cross analyzed by TRF performed on the spore progenies (see also **Figure 5C**). **(D)** TRF Southern blots of 3 insertion mutants from the CliP library in genes not annotated to be telomere-related compared to the parent CC4533- strain. Mutant A = LMJ.RY0402.203280; mutant B = LMJ.RY0402.060361; mutant C = LMJ.RY0402.166033. **(E)** Telomerase-negative spore progenies A1-4 and B1-4 are shown after about six months of maintenance on solid media. A4 and B1 experienced growth defect and cell death typical of replicative senescence but generated clonal low-frequency survivors after additional restreaks. The telomerase-negative spores A1 and B2 did not show senescence phenotype. Right: summary table of the presence (red) or absence (green) of senescence phenotype in telomerase-negative (paromomycin-resistant) progenies of tetrads from the indicated crosses.

**Supplemental Table S1: Primers used for telomere-PCR, PETRA, hairpin assay and PCR verification of CliP mutants**

| **Name** | **Sequence (5'→3')** | **Description** |
|---|---|---|
| oT1090 | ACAAAACCCCTTAAAACCCCCATTTAG | Forward primer for Telomere-PCR |
| 169M | CGGGATCCGGGGGGGGGG | Poly-G Reverse primer for Telomere-PCR |
| oT0958 (CHSB from (Fulneckova et al., 2013) | GTTTTAGGGTTTTAGGGTTTTAGGGTTTTAG | Telomere-specific probe for TRF Southern blot |
| oT1053 | TGTGTGTGTGTGTGTGTGTGTGTGTG | Used in TRF Southern blot to check for TG |

| | | |
|---|---|---|
| | | microsatellites |
| $F_{m1}$ | AACACATTTGCACACGCAAT | Forward primer used to identify insertion in tel-m1 as suggested in https://www.chlamylibrary.org/feature/LMJ.RY0402.077111_2 |
| $R_{m1}$ | GGAGTGGGGCACAAAGTAGA | Reverse oligo used to identify insertion in tel-m1 as suggested in https://www.chlamylibrary.org/feature/LMJ.RY0402.077111_2 |
| $F_{m2}$ (F12) | TCTGGATCCAAATCCACCGC | Forward primer used to characterize tel-m2 by the absence of a band, present in CC4533- and tel-m1. |
| $R_{m2}$ (R6) | GACCCTGCCACTGCCTTATT | Reverse primer used to characterize tel-m2 by the absence of a band, present in CC4533- and tel-m1. |
| $R_{m4}$ | GGAGATAGCCTGTGAGCCAG | Primer used for the characterization of tel-m4 with primer $K_2$. https://www.chlamylibrary.org/feature/LMJ.RY0402.105594_2 |
| $K_1$ (oMJ913) | GCACCAATCATGTCAAGCCT | 5' primer relative to the paromomycin cassette used in the generation of the CliP library (https://www.chlamylibrary.org/content/Update-how-characterize-insertion-sites-PCR-different-primers-are-needed-mutants-starting) |
| $K_2$ (oMJ944) | GACGTTACAGCACACCCTTG | 3' primer relative to the paromomycin cassette used in the generation of the CliP library |

| | | |
|---|---|---|
| | | (https://www.chlamylibrary.org/content/Update-how-characterize-insertion-sites-PCR-different-primers-are-needed-mutants-starting) |
| oT1208 ($F_c$; F2) | GCAAACGCTTCATCAGGCAA | Forward primer in the telomerase gene not affected in the WT nor the telomerase mutants |
| oT1209 ($R_c$; R2) | GCGTATGACCTACCGGCTAC | Reverse primer in the telomerase gene not affected in the WT nor the telomerase mutants |
| OLIP1 | CCGCACATGAGACGTTACAG | Determination of mating-type + of Chlamydomonas strains. |
| OLIP2 | GATTGCTCTGTCGTTGCAGA | Determination of mating-type + of Chlamydomonas strains. |
| OLIM1 | TGGCGTACCTTTCTGTAGGG | Determination of mating-type - of Chlamydomonas strains. |
| OLIM2 | GCCACG AAG GCAGTTACATT | Determination of mating-type - of Chlamydomonas strains. |
| Blunt Hairpin | GGATCCGACTTTTGTCGGATCC | Used in the hairpin assay. |
| PETRA-T | CTCTAGACTGTGAGACTTGGACTACCCTAAAACCCT | Used in the PETRA experiment. |
| PETRA-A | CTCTAGACTGTGAGACTTGGACTAC | Used in the PETRA experiment. |
| 1R(a) | TACTTGTGTGTGCTGTGCGT | Used in the PETRA experiment for chromosome 1R. |
| 9R(a) | ACAGCACAATACAGTATATA | Used in the PETRA experiment for chromosome 9R. |
| 10R(c) | AACGTCCTCGTGAGACCACC | Used in the PETRA experiment for chromosome 10R. |
| Cr18S_PR1_F | CTTCACTGTCTGGGACTCGGA | Forward primer to amplify a |

| | | |
|---|---|---|
| | | fragment in 18S ribosomal subunit gene from genomic DNA of strain CC4350 |
| Cr18S_PR1_R | ACTAAGAACGGCCATGCACCA | Reverse primer to amplify a fragment in 18S ribosomal subunit gene from genomic DNA of strain CC4350 |

#### SUPPLEMENTAL METHODS

##### Strains and growth conditions

Strains T222+, S24-, CC124-, CC125+, CC503+, CC1009-, CC1010+ and CC4425+ (D66) are described in (Gallaher et al., 2015). Strain 21gr is described in (Harris, 2009). Strains cw15.J3- and cw.J14- are cell-wall-less strains obtained by crossing. Strains CC620+, CC521+ and CC4350+ are described in the Chlamydomonas Resource Center (https://www.chlamycollection.org/). Strain 4533- is described in (Li et al., 2016). Strain S1D2 is described in (Gross et al., 1988; Harris, 2001; Vysotskaia et al., 2001). Unless stated otherwise cells were grown under continuous illumination either on plates or in agitated 200 mL liquid cultures in Tris-Acetate-Phosphate (TAP) medium (Harris, 2009) under low-light (LL), *i.e.* 8 $\mu E.m^{-2}.s^{-1}$ or higher-light (HL), i.e. 80 $\mu E.m^{2}.s^{-1}$.

##### gDNA extraction

Unless stated otherwise, cells were grown in liquid cultures to early stationary phase (~$2.10^{7}$ $cells.mL^{-1}$) and 150 mL were collected by centrifugation (5000 g, 5 min). The pellet was frozen at -80°C. Cells were then thawed at room temperature and 5 mL of preheated buffer AP1 with RNAse (Qiagen DNA Plant Maxi Kit) was added and cells lysed at 65°C for 2 hours. After lysis, gDNA was extracted according to the manufacturer's protocol (Qiagen DNA Plant Maxi Kit).

##### Telomere PCR and sequencing

Bulk genomic DNA was denatured at 95°C during 5 min. End-labeling reactions (total volume 6 μL) contained 100 ng of bulk genomic DNA, 1X New England Biolabs Restriction Buffer 4, dCTP 100 μM and 1 unit of Terminal Transferase (New England Biolabs, NEB) and was carried out at 37°C during 30 min then 65°C during 10 min and 94°C during 5 min. The end-labeled telomeres were then amplified with the primers 169M (poly-G containing primer) and oT1090 targeting the subtelomere/telomere junction common to 10 telomeres of 8 chromosomes. PCR reactions (40 µL) contained the end-labeled DNA, 200 µM of dNTPs, primers at 0.5 µM for oT1090 and 0.75 µM for 169M, 1X Taq Mg-free Buffer (NEB) and 2.5 U of Standard Taq Polymerase (NEB). The PCR conditions were: 94°C 3 min; 32 cycles of 94°C 20 sec, 60°C 40 sec, 68°C 20 sec; 68°C 5 min.

For sequencing, PCR products were ligated for 1h at 16°C in a pDrive plasmid. 2 µL of the ligation product was transformed into competent bacteria (PCR cloning kit, Qiagen). Bacteria were plated on LB + Ampicillin (100 µg/mL) + IPTG (50 µM) + X-gal (80 µg/mL) medium over night at 37°C. Plasmids were extracted and purified (Millipore Plasmid Miniprep 96 Kit and Manifold) after 24 h of culture of white colonies in 1 mL of LB 2X + Ampicillin (100 µg/mL) in a 96-well microplate. DNA insertion in plasmids was verified by *Eco*RI (NEB) digestion. Plasmids were Sanger sequenced with M13-PU primer (Eurofins Genomics).

**Isolation of nuclei**

Nuclear fraction was prepared from cell wall mutant CC4350. Exponentially growing cells (2 days in liquid culture) were gently spun and thoroughly resuspended in 90 mL Buffer A per liter of culture (25 mM Hepes-NaOH, pH 7.5, 20 mM KCl, 20 mM $MgCl_2$, 600 mM sucrose, 10% glycerol, 5 mM DTT). Triton X-100 was first diluted in 10 mL of Buffer A per liter of culture and subsequently added drop-wise to the cells while swirling them gently, to a final concentration of 0.5%. Nuclei were pelleted at 800 g for 2-4 minutes. Using a paintbrush, pellet was gently resuspended in fresh Buffer A without Triton X-100. After centrifugation, integrity of nuclei (1-5 µL) was checked by fluorescent microscopy using DAPI/vectashield (5 µL) staining. Nuclei were resuspended in Buffer B (2.5% Ficoll 400, 0.5 M sorbitol, 0.008% spermidine, 50% glycerol, 1 mM DTT), using 1-2 mL per 200 mL of original culture volume. For storage, nuclei were frozen in liquid nitrogen and stored at -80°C.

**Micrococcal nuclease hypersensitivity assay**

Micrococcal nuclease hypersensitivity assay was based on (Lodha and Schroda, 2005). One milliliter of nuclei isolated from *C. reinhardtii* strain CC-4350 cw15 mt+ in Buffer B was thawed on ice. To collect nuclei, sample was spun at maximal speed for 15 seconds and resuspended in 500 µL of 1x MN Buffer (50 mM Tris-HCl pH 8.0, 5 mM $CaCl_2$). Reactions of total volume of 110 µL were carried out in 1x MN Buffer using 60 µL of sample and different amounts of micrococcal nuclease units (Fermentas). Genomic DNA from *C.*

*reinhardtii* CC-4350 (~750 ng) was used as a control for enzyme activity and digested with 15 U of nuclease. Samples were incubated 3 minutes at room temperature and reactions were stopped by adding 110 μL of STOP buffer (1% SDS, 50 mM EDTA). Proteins were then denaturated for 45 min at 65°C and DNA was extracted using 500 μL of phenol:chloroform:isoamyl alcohol (25:24:1) using phasetrap A (Peqlab). Aqueous phase was precipitated by adding 42 μL of 3 M NaOAc and 840 μL of 96% ethanol. DNA was pelleted by centrifugation for 10 min at maximal speed. Pellet was washed with 70% ethanol, dried and resuspended in 25 μL $H_2O$. 6x loading dye (6 μL) was added prior loading onto 1.5% agarose gel. DNA was stained with ethidium bromide (1% solution, AppliChem), blotted onto uncharged membrane (Amersham) and hybridized with a $(T_3AG_3)_3$ probe. After scanning, membrane was stripped and reprobed using 18S-derived probe.

**PETRA and hairpin assay**

Telomere length of individual chromosomes was determined by Primer Extension Telomere Repeat Analysis (PETRA) as previously described (Watson et al., 2016). For primer extension, we used the *C. reinhardtii* specific PETRA-T oligonucleotide 5′-CTCTAGACTGTGAGACTTGGACTACCCTAAAACCCT-3′. For specific chromosome arms we used subtelomeric oligonucleotides 1R: 5′-TACTTGTGTGTGCTGTGCGT-3′, 9R: 5′-ACAGCACAATACAGTATATA-3′ and 10R: 5′-AACGTCCTCGTGAGACCACC-3′. The hairpin assay for detecting blunt-ended telomeres was performed as previously described (Kazda et al., 2012). Southern hybridization was done with a [$^{32}$P]ATP-labeled $(TTTTAGGG)_4$ probe. PETRA membrane was also hybridized with [$^{32}$P]ATP-labeled 1 kb ladder (Thermo Scientific).

**TRF Southern blot analysis**

2 μg of genomic DNA was digested in 300 μL with a cocktail of 6 restriction enzymes (*Pst*I, *Bam*HI, *Mnl*I, *Fok*I, *Taq*I and *Msp*I; 20 units each). Digestion products were isopropanol precipitated, resuspended in loading buffer (Gel Loading Dye, Purple 6X, New England BioLabs) and resolved on a 1.5% agarose gel for 4h at 150V. The gel was then soaked in a denaturation bath (0.4 M NaOH and 1 M NaCl) for 20 min and transferred overnight by capillarity to a charged nylon membrane (Hybond XL, GE Healthcare). The CHSB *Chlamydomonas* telomere-specific oligonucleotide probe (Fulneckova et al., 2013) (5′-GTTTTAGGGTTTTAGGGTTTTAGGGTTTTAG-3′) was $^{32}$P-labelled at the 5′ terminus

with ATP (γ-$^{32}$P) by the T4 polynucleotide kinase (New England BioLabs). The membrane was hybridized using the Rapid-hyb buffer protocol (GE Healthcare). In brief, the membrane was pre-hybridized at 42°C in Rapid-hyb buffer for 1 h, then the radioactive probe (20 pmol) was added and the incubation was continued for 1 h. The membrane was washed consecutively with 5X SSC, 0.5% SDS (42°C for 10 min); 5X SSC, 0.1% SDS (42°C for 20 min); and 1X SSC, 0.1% SDS (25°C for 30 min). The membrane was then imaged with a Typhoon FLA 9500 scanner (GE Healthcare). Average telomere length was assessed using ImageJ 1.49v (NIH) by measuring the peak of the telomere length distribution signal.

### Supplemental Figure S1

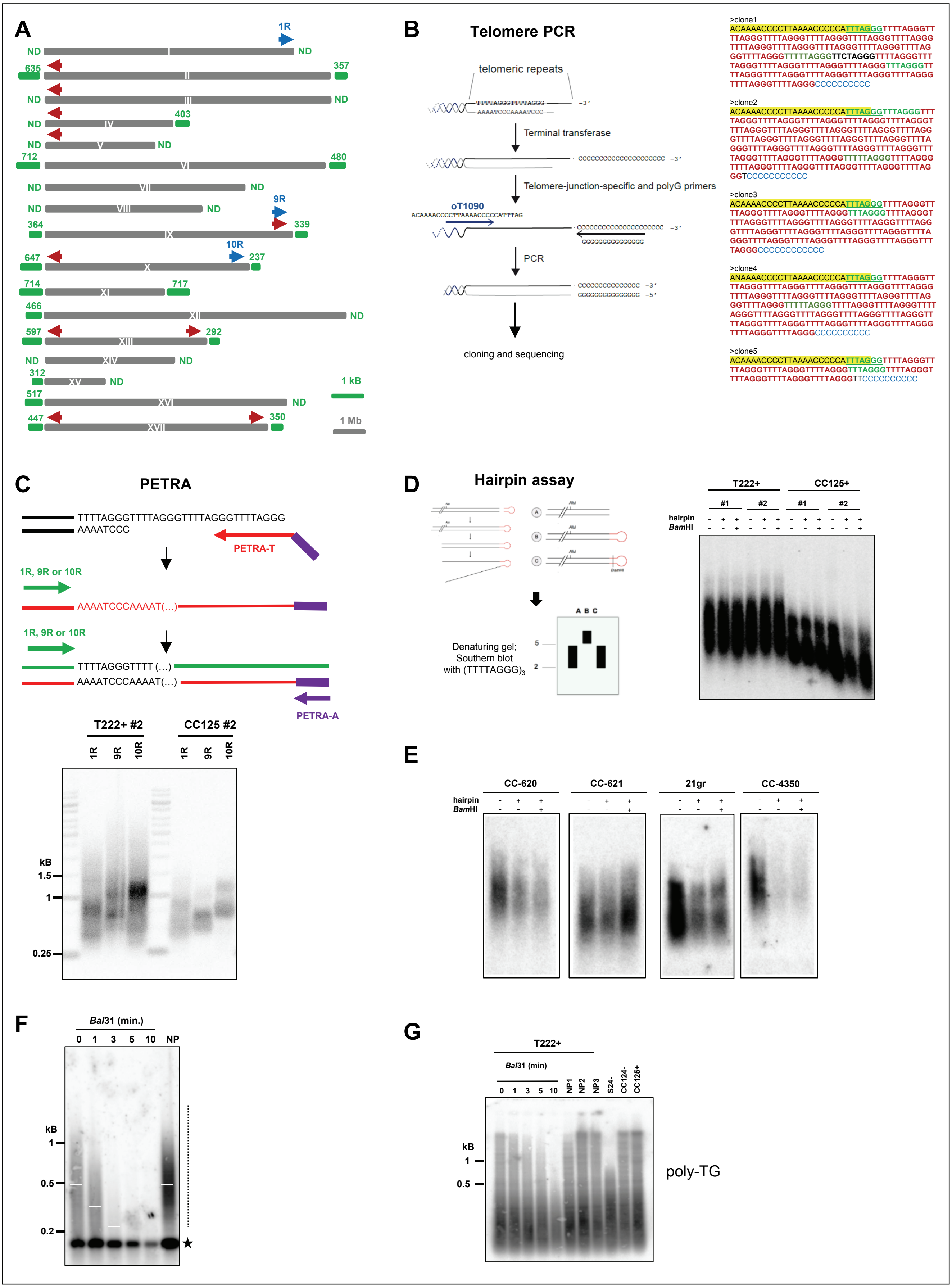

**Supplemental Figure S1: Characterization of *C. reinhardtii* telomere repeats by telomere-PCR and PETRA. Related to Figure 1. (A)** Scheme of the 17 *C. reinhardtii* chromosomes. Red arrow: forward primer oT1090 used in telomere-PCR. Blue arrows: forward primers 1R, 9R and 10R used in PETRA experiments. Green numbers refer to telomere length from the public genome sequence of *C. reinhardtii* (Phytozome genome version 5.5; https://phytozome.jgi.doe.gov/pz/#!info?alias=Org_Creinhardtii). ND: no telomere in the database. **(B)** Scheme of the different steps of telomere-PCR (left). After adding a poly-C extension to telomeric ends by terminal transferase, PCR amplification using a forward primer at a subtelomere-telomere junction (oT1090) and a poly-G reverse primer was performed. The resulting PCR products were cloned and sequenced. Telomere-PCR was used to amplify telomeric repeats from the T222+ strain. The resulting PCR products were cloned in *E. coli* and 32 clones sequenced. Five examples of telomere-PCR sequences showing the sequence of primer oT1090 (yellow), the poly-C extension (blue), canonical *C. reinhardtii* repeats (TTTTAGGG, in red) and variant motifs (other colors) (right). The frequency of the observed repeats is indicated in **Table 1**. **(C)** Scheme of the PETRA assay for 3′ overhang detection. The primer PETRA-T was annealed to the putative 3′ overhang and extended using DNA PolI. PCR amplification was then performed using a subtelomere-specific primer and PETRA-A primer, which annealed to the non-telomeric sequence of the 5′ end of PETRA-T. (top). An independent biological replicate of the result shown in **Figure 1B** is shown at the bottom. **(D)** Left: scheme of the hairpin assay to detect blunt-end telomeres as described (Kazda et al., 2012). Right: two independent biological replicates for strains T222+ and CC125+ show no evidence for blunt telomeres. **(E)** Four other *C. reinhardtii* strains CC-620, CC-621, 21gr and 302, were tested and were negative for blunt ends by hairpin assay. **(F)** Independent biological replicate of the *Bal*31 assay performed on strain T222+ (see **Figure 1D**). **(G)** A poly-TG probe did not detect the band observed at ~200 bp with a telomere probe. Genomic DNAs from strains T222+, S24-, CC124- and CC125+ were digested with the same protocol used for TRF experiments, Southern blotted and then probed with a radioactive $(TG)_{13}$ probe. T222+ samples were also treated by *Bal*31 exonuclease for increasing amounts of time (5 first lanes) before digestion with the restriction enzymes. NP1-3 samples were not column-purified.

### Supplemental Figure S2

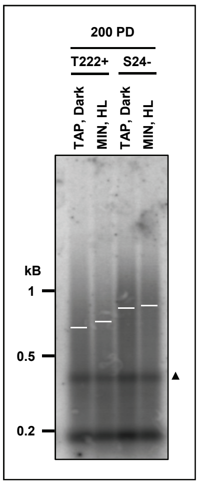

**Supplemental Figure S2: Prolonged culture in different growth conditions does not significantly affect telomere lengths. Related to Figure 2.** Reference strains T222+ and S24- were grown in either heterotrophic conditions (TAP, Dark) or in photo-autotrophic conditions (MIN, HL). When reaching the stationary phase, cultures were diluted in fresh media and this was repeated for 2 months, *i.e.* ~200 population doublings (PD) before TRF analysis. Triangle: band not seen in other TRF experiments.

### Supplemental Figure S3

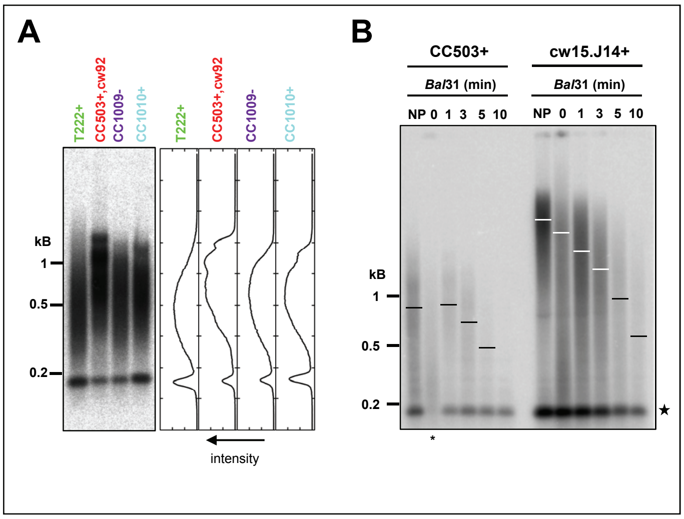

**Supplemental Figure S3: Multimodal and very long telomeres from CC503+ and cw15.J14+ correspond to terminal DNA fragments. Related to Figure 3. (A)** TRF Southern blot of biological replicates of the indicated strains shown in **Figure 3** (left), with intensity profile analysis (right) showing the multimodal distribution of CC503+ and CC1010+. **(B)** Genomic DNAs of strains CC503+ and cw15.J14+ were subjected to *Bal*31 exonuclease digestion for 1 to 10 minutes prior to digestion by the restriction enzyme cocktail and Southern blotting using the CHSB (oT0958) probe, showing that the detected signal is indeed *Bal*31-sensitive and hence represents terminal fragments.

### Supplemental Figure S4

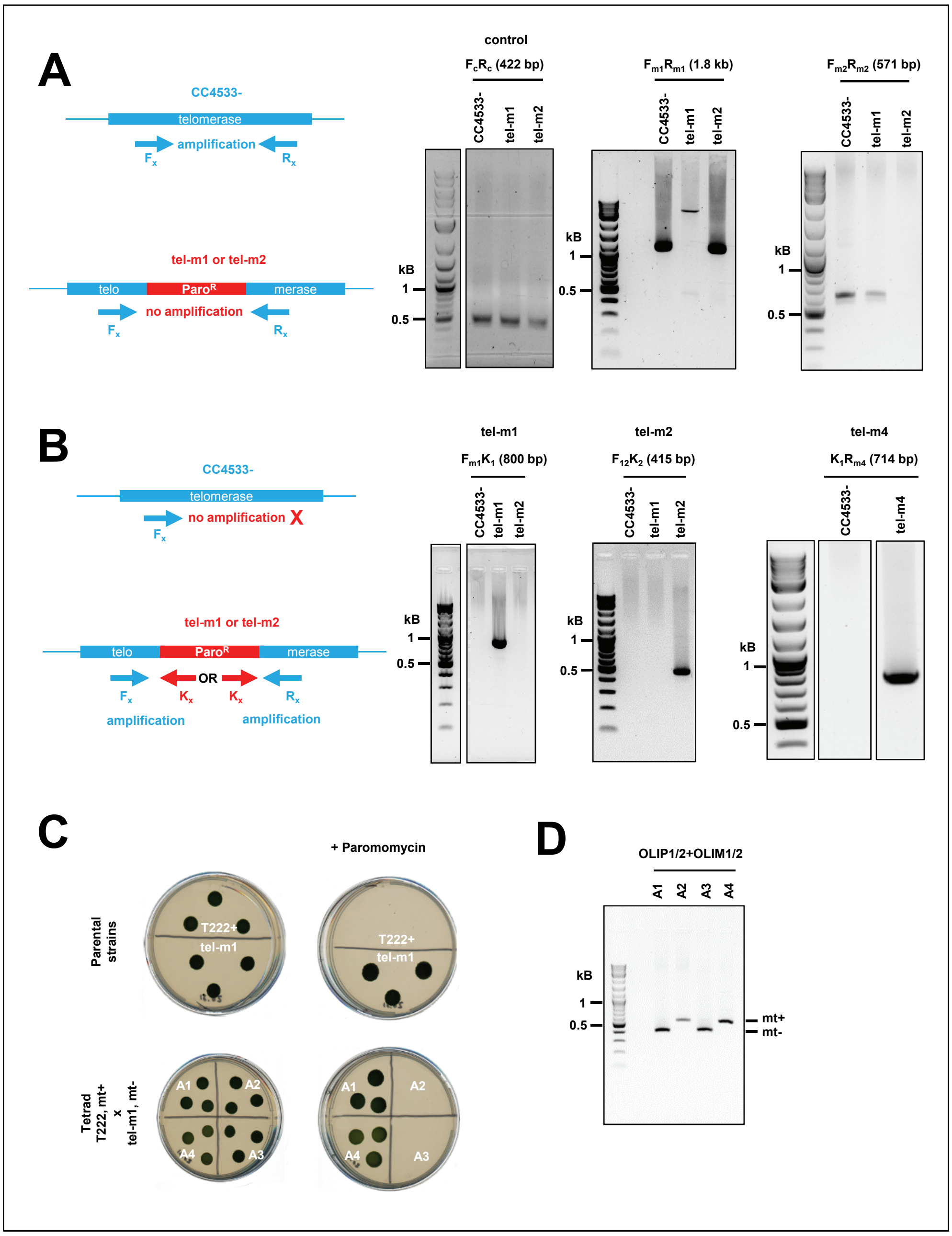

**Supplemental Figure S4: PCR verification of insertion mutants tel-m1 and tel-m2. Related to Figure 4. (A)** Scheme of expected PCR amplification product (left). $F_x$ and $R_x$ denote primers where "x" stands for "m1" or "m2". PCR using primers specific to the reported insertions sites in tel-m1 ($F_{m1}$ and $R_{m1}$) and tel-m2 ($F_{m2}$ and $R_{m2}$) gave no signal in the targeted mutant because the inserted marked is loo large for efficient amplification, but amplified the expected product in the reference strain and the other mutant (of size 1.8 kb and 571 bp, respectively; gels in the middle and on the right). Primers $F_c$ and $R_c$ amplified of a 422-bp product in the *CrTERT* gene outside of the predicted insertion sites in tel-m1 and tel-m2 (positive control, gel on the left). **(B)** Scheme of expected PCR amplification product using specific primers for tel-m1 ($F_{m1}$) and tel-m2 ($F_{12}$), combined with primers in the paromomycin cassette ($K_1$ and $K_2$) (left). The two pairs of primers amplified the expected products in the respective mutants (right). All bands were excised, gel-purified and sequenced**.** PCR-amplification using primers $K_1$ and $R_{m4}$ were used to characterize the insertion in tel-m4. The specific band obtained in tel-m4 was excised from the gel and sequenced and could indeed be mapped to the expected *CrTERT* sequence. **(C)** Mutant tel-m1 was crossed with T222+ strain. The cross with tel-m2 is not shown. The parental strains (tel-m1 and T222+) as well as the progeny of the four spores of a tetrad from the diploid are spotted on TAP solid media with ("+ Paromomycin") or without paromomycin (10 µg.mL$^{-1}$). Five tetrads were analyzed for each cross for the segregation of paromomycin resistance. **(D)** The 2:2 segregation of the mating type was checked by PCR.

### Supplemental Figure S5

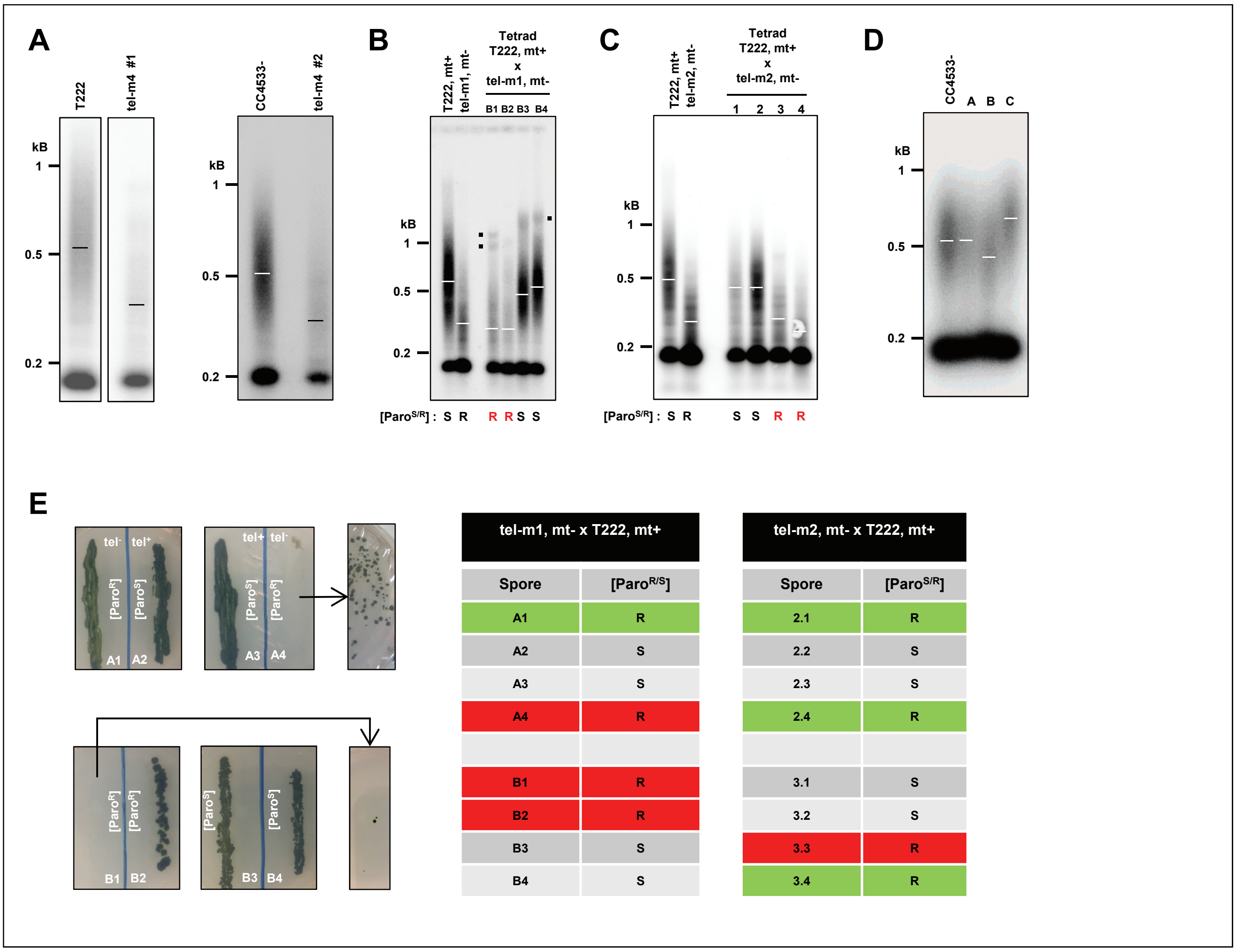

| tel-m1, mt- x T222, mt+ | |
|---|---|
| Spore | [Paro$^{R/S}$] |
| A1 | R |
| A2 | S |
| A3 | S |
| A4 | R |
| B1 | R |
| B2 | R |
| B3 | S |
| B4 | S |

| tel-m2, mt- x T222, mt+ | |
|---|---|
| Spore | [Paro$^{S/R}$] |
| 2.1 | R |
| 2.2 | S |
| 2.3 | S |
| 2.4 | R |
| 3.1 | S |
| 3.2 | S |
| 3.3 | R |
| 3.4 | R |

**Supplemental Figure S5: Telomere length and senescence phenotype of *CrTERT* mutants. Related to Figure 5. (A)** Insertion mutant tel-m4 has shorter telomeres than T222+ and CC4533- as shown by TRF Southern blots of two biological replicates of the tel-m4 mutant strain. **(B)** Independent tetrad of the tel-m1 x T222 cross analyzed by TRF Southern performed on the spore progenies (see also **Figure 5B**). **(C)** Independent tetrad of the tel-m2 x T222 cross analyzed by TRF performed on the spore progenies (see also **Figure 5C**). **(D)** TRF Southern blots of 3 insertion mutants from the CliP library in genes not annotated to be telomere-related compared to the parent CC4533- strain. Mutant A = LMJ.RY0402.203280; mutant B = LMJ.RY0402.060361; mutant C = LMJ.RY0402.166033. **(E)** Telomerase-negative spore progenies A1-4 and B1-4 are shown after about six months of maintenance on solid media. A4 and B1 experienced growth defect and cell death typical of replicative senescence but generated clonal low-frequency survivors after additional restreaks. The telomerase-negative spores A1 and B2 did not show senescence phenotype. Right: summary table of the presence (red) or absence (green) of senescence phenotype in telomerase-negative (paromomycin-resistant) progenies of tetrads from the indicated crosses.
